# The MAP2Ks can use ADP to phosphorylate and activate their substrate MAPKs *in vitro*

**DOI:** 10.1101/2024.02.22.581546

**Authors:** Pauline Juyoux, Jill von Velsen, Erika Pellegrini, Matthew W. Bowler

## Abstract

Kinases are a diverse group of enzymes that use ATP to phosphorylate a variety of substrates. Protein kinases evolved in eukaryotes as important mediators of cell signalling that target specific amino acid side chains to modulate downstream protein function. Among them, the MAPKs (mitogen-activated protein kinases) are a family of intracellular protein kinases that form signalling cascades responding to a number of stimuli, that control fundamental mechanisms such as proliferation, differentiation, inflammation and cell death. Signals propagate through consecutive kinases which eventually phosphorylate and activate a MAPK. Here, we show that the dual specificity threonine/tyrosine MAP kinase kinases (MAP2Ks or MEKs) are able to phosphorylate and activate their substrate MAPKs using ADP as well as ATP *in vitro*. As the pathways are involved in the stress response, we speculate that it would represent an advantage to be able to maintain signalling under conditions such as hypoxia, that occur under a number of cell stresses, including cancer and atherosclerosis, where the available pool of ATP could be depleted.

## Introduction

Phosphorylation is a post-translational modification that is extensively used in the control of processes in higher organisms (1). Protein kinases are important mediators of cell signalling by targeting specific amino acids for phosphorylation, which modulates downstream protein function. One such family of protein kinases is the MAPKs (mitogen-activated protein kinases) that form signalling cascades, responding to a number of stimuli, that control fundamental mechanisms such as proliferation, differentiation, inflammation and cell death (2). Signals propagate through kinases which eventually phosphorylate and activate a MAPK, through double phosphorylation at a TxY motif in the activation loop (A-loop) by a specific MAPK kinase (MAP2K). Once activated, the MAPK modulates the expression of genes through activation of transcription factors or other protein kinases (3, 4).

Kinases have evolved to phosphorylate their substrates using ATP as it is the ubiquitous currency for energy transduction in cells and is around 10 to 1000 times more abundant than ADP in the cell(5). ADP dependent kinases were initially identified in extremophiles (6-8) and were originally thought to be an adaptation to high temperatures, or be an evolutionary remnant of an ancient metabolic pathway. Subsequently, ADP dependent kinases have been identified in all kingdoms of life, including mesophiles (9) and vertebrates (10-13). However, they are rare, cannot use ATP, are restricted to sugar and cysteine substrates (14) and an ADP dependent protein kinase has never been identified. Nucleophilic attack on the β-phosphate of ADP is chemically similar to the γ-phosphate of ATP, with the same standard transformed Gibbs energies of formation in physiological conditions (15). However, as most enzymes have evolved to precisely position substrates for nucleophilic attack, meaning they are either ATP or ADP dependent, there are no examples of enzymes that can use both nucleotides to phosphorylate a substrate. Additionally, the concentration of ATP is usually much higher than ADP making it a more abundant substrate, with the distance from equilibrium driving the many processes that depend on ATP. Many proteins are sensitive to this ratio, such as ATP dependent potassium channels (16) and the pseudo kinase domain of IRE1 (17), moving from an ATP bound to an ADP bound state, allowing cells to react to the metabolic state. This is important as there are many conditions that lead to drastic changes in the ATP/ADP ratio, such as hypoxia, that require cells to react. Here, we demonstrate that the human MAP2K dual specificity protein kinases can use both ATP and ADP to transfer a phosphate group to the A-loop of their target MAP kinases *in vitro*.

### The activation loop of p38α is phosphorylated on residues Thr180 and Tyr182 by MKK6^DD^ in the presence of ADP

The active conformation of MKK6, as for the other MAP2Ks, is stabilised by the double phosphorylation of serine and threonine residues in their A-loops (S207 and T211 in MKK6). We used a constitutively active mutant, named MKK6^DD^, where these two residues are mutated to aspartates to mimic phosphorylation. While preparing complexes for structural studies (18), we assessed the ability of MKK6^DD^ to phosphorylate p38α in the presence of different nucleotides. Native PAGE gels were used to separate the different phosphorylation states of p38α (Figure 1A) with the bands identified by mass spectrometry. As expected, in the presence of ATP, MKK6^DD^ transfers two phosphate groups to p38α (p38α-2P) (and also, to a lesser degree, a third (p38α-3P)), indicating high activity. We also observed that in the presence of ADP, MKK6^DD^ can phosphorylate p38α, resulting in a mixture of both mono- (p38α-1P) and bi-phosphorylated (p38α-2P) p38α. We obtained similar results using MKK6^WT^, which displays basal activity, and using p38α^K53R^ (a p38α kinase-dead mutant) (Figure S1A), ruling out p38α auto-phosphorylation. We complemented this analysis with native ESI-QTOF mass spectrometry experiments, as well as LC-MS/MS detection of phosphorylated sites on samples in solution (Figure 1B and C), that confirmed that both the T180 and Y182 residues of the p38α A-loop were phosphorylated by MKK6^DD^ in the presence of either ADP or ATP with no preference for one residue over the other. In order to rule out ATP contamination we repeated the experiments with ultra pure ADP, obtaining the same result, and confirmed the absence of ATP in our ADP stocks via natural abundance ^31^P NMR (Figure S2).

**Figure 1.**
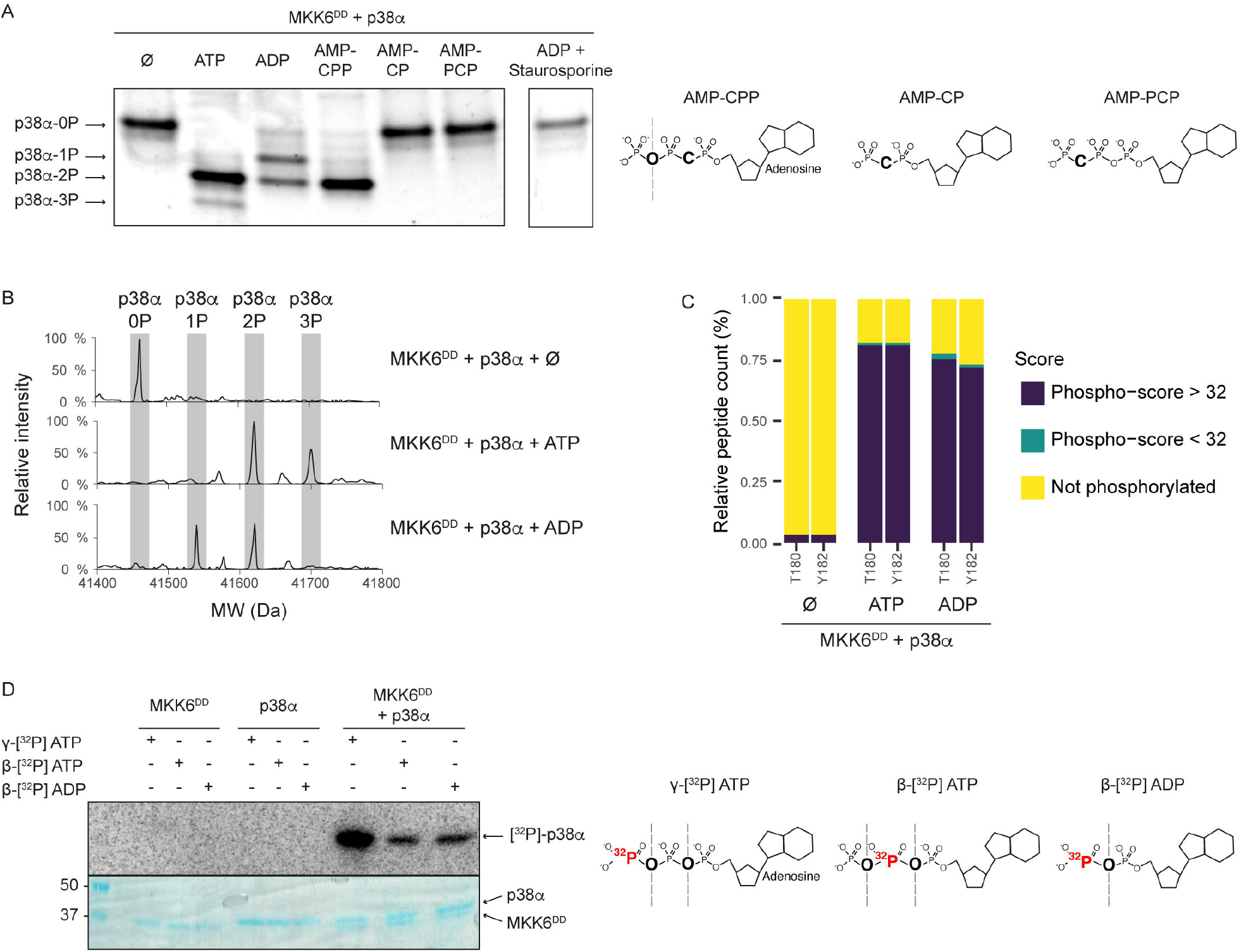
MKK6 can phosphorylate p38α using ADP *in vitro*. (A) Native PAGE gel of MKK6^DD^ + p38α in the presence of different nucleotides and an inhibitor. The p38α bands run depending on the number of phosphorylated residues. Chemical structures of the different nucleotide analogues are represented. The dashed lines represent bonds that are available for hydrolysis by the kinase. AMP-CPP = Adenosine-5’-[(α,β)-methyleno]triphosphate, AMP-CP = Adenosine-5’-[(α,β)-methyleno]diphosphate and AMP-PCP = Adenosine-5’-[(β,γ)-methyleno]triphosphate. See Fig S1 for the uncropped gel. (B) Measurement by ESI-QTOF MS of the phosphorylation of p38α by MKK6^DD^. Each phosphorylation increases the mass by 80 Da. (C) Phosphorylation of the p38α A-loop T180 and Y182 residues by MKK6^DD^ measured by LC-MS/MS. Results are represented as the percentage of peptide count detected as not phosphorylated (yellow), potentially phosphorylated (blue) and phosphorylated (purple) by MASCOT analysis. (D) *In vitro* phosphorylation assay with radiolabelled nucleotides. Samples of MKK6^DD^ and p38α were incubated with γ-[^32^P] ATP, β-[^32^P] ATP, or β-[^32^P] ADP. Samples were separated by SDS-PAGE and visualized by autoradiography (top) and Coomassie stain (bottom). Chemical structures of the different radiolabelled nucleotides are represented, with the radiolabelled ^32^P in red and bonds available for hydrolysis are indicated with dashed lines.

### MKK6^DD^ can use the β-phosphate of both ADP and ATP to phosphorylate p38α

Direct proof of the incorporation of the β-phosphate of ADP was obtained with a radionucleotide assay (Figure 1D). Incubation of MKK6^DD^ and p38α in the presence of either β-[^32^P] ADP or β-[^32^P] ATP resulted in the transfer of the labelled phosphate to the substrate. Given that the β-phosphate can be transferred from either ATP or ADP we initially speculated that the dual phosphorylation could proceed via a single step, using first the γ-phosphate of ATP and, subsequently, the β-phosphate of the resulting ADP product in a fully processive mechanism. However, by incubating MKK6^DD^ and p38α with AMP-CPP (Adenosine-5’-[(α,β)-methyleno]triphosphate), an ATP analogue from which the β-phosphate cannot be cleaved, we demonstrated the use of the β-phosphate is not required to achieve dual phosphorylation (Figure 1A) and ADP phosphorylation is therefore an alternative route to phosphorylation of the A-loop. Additional controls using the non-cleavable nucleotide analogues AMP-CP (Adenosine-5’-[(α,β)-methyleno]diphosphate) and AMP-PCP (Adenosine-5’-[(β,γ)-methyleno]triphosphate), as well as the kinase competitive inhibitor staurosporine, showed no phosphorylation activity, demonstrating that the transfer of phosphate in the presence of ADP occurs via nucleophilic attack of the β-phosphate in the active site of MKK6.

### Phosphorylation with ADP is less efficient than with ATP

We then proceeded to a time course phosphorylation assay with either ADP or ATP (Figure 2) showing that phosphorylation using ADP is ~100 times slower than with ATP, but the curves are similar between the nucleotides and match previous studies with ATP (19, 20). The point at which half of the p38α is phosphorylated at least once, under the conditions used for our assay, is reached in 21 seconds with ATP and 2243 seconds (37 minutes) with ADP. While this is significantly slower, it is still on a biologically relevant timescale.

**Figure 2.**
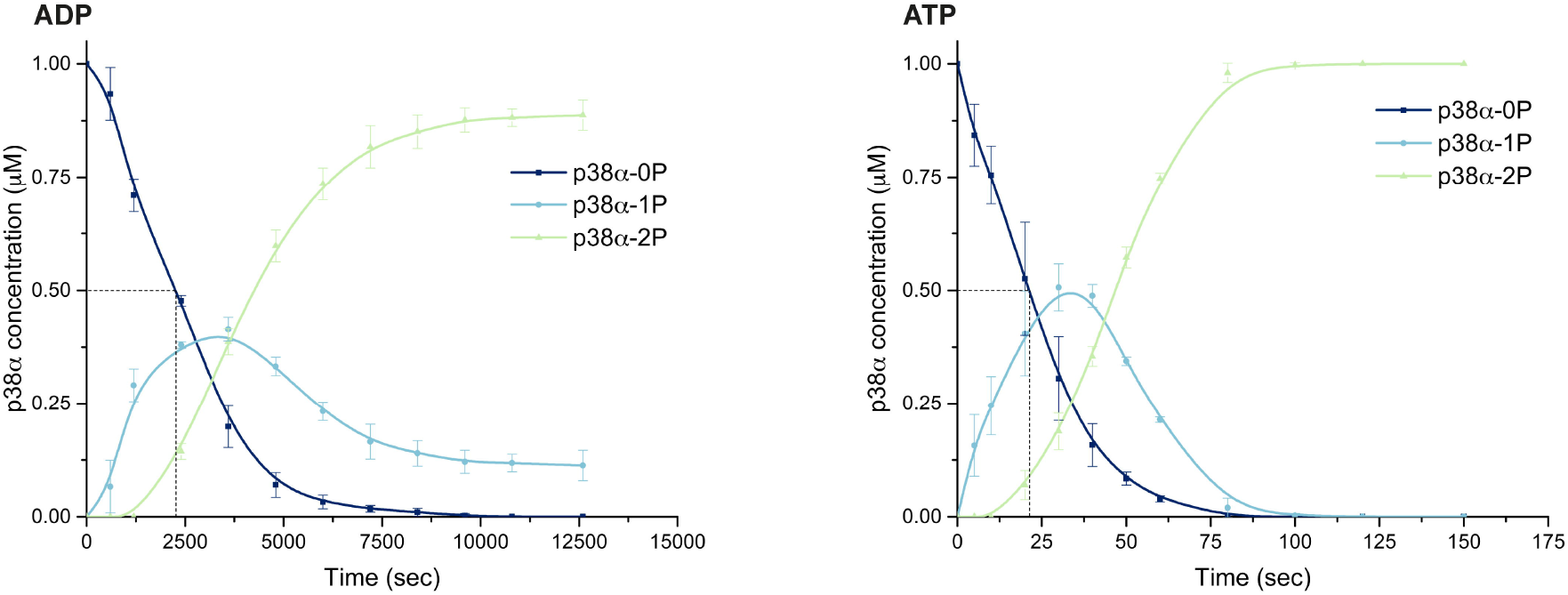
Phosphorylation using ADP is slower than with ATP. Time course phosphorylation assay of p38α by MKK6^DD^ with ADP (left) and ATP (right). The dashed line represents the point at which half of the p38α population is phosphorylated at least once, 2243 seconds (37 minutes) with ADP and 21 seconds with ATP. Concentrations of the different phosphorylation states of p38α were calculated for triplicates of native PAGE gels and fit with a b-spline curve. See Fig S1 for representative gels.

The *in vitro* dual phosphorylation of p38α by MKK6^DD^, in the presence of both ATP and ADP, appears to occur through a distributive mechanism, with first a pool of mono-phosphorylated p38α population building up, and the bi-phosphorylated population appearing afterwards (Figure 2). This delay in the formation of bi-phosphorylated p38α was previously observed (19), and is indicative of a distributive mechanism as a single first product is produced quickly before re-binding leads to the second phosphorylation event.

Attempts to perform a complete kinetic study of the system faced limitations. Given the difference in speed between the reaction using ADP or ATP, it was difficult to find experimental conditions (temperature, kinase and substrate concentration, time course) measurable and comparable with both nucleotides. In addition, the dual phosphorylation of the p38α A-loop by MKK6 does not follow classical Michaelis-Menten kinetics. As predicted by simulation studies and assays with the MEK1-ERK2 system (21), a non-processive (or distributive) mechanism predicts that there will be a paradoxical decrease in the rate of MAPK phosphorylation as the MAPK concentration is increased, a phenomenon we also observed.

In some experimental conditions, the dual phosphorylation of p38α with ADP was incomplete during the experimental timeline (Figure 1A and S1). Kinetic studies with ATP have shown that the second phosphorylation reaction occurs more slowly than the first (19, 20). This would be enhanced in the slower reaction with ADP, with the phospho-group of p38α-1P making it more difficult for the second phospho-acceptor to access the ADP, possibly due to steric hindrance.

### All MAP2Ks can use ADP to phosphorylate their MAPK substrates

To understand the extent of the ability of protein kinases to use ADP to phosphorylate substrates, we tested other activated MAP2K–MAPK pairs. All the tested MAP2Ks were able to phosphorylate their target MAPKs using either ATP or ADP (Figure 3), with different relative activities depending on the MAP2K. In comparison to the reduced MKK6 activity with ADP, MKK4 appears to be similarly active on p38α with either ATP or ADP (Figure 3C). MKK4 and MKK7 activity on JNK1 was similar with either ATP or ADP as a phosphate source. MKK7 might even be slightly more active with ADP than with ATP and also autophosphorylates using ADP (Figure 3E). The activity of both MEK1 and MEK2 on ERK1 was low with ADP in comparison to ATP, but still occurs.

**Figure 3.**
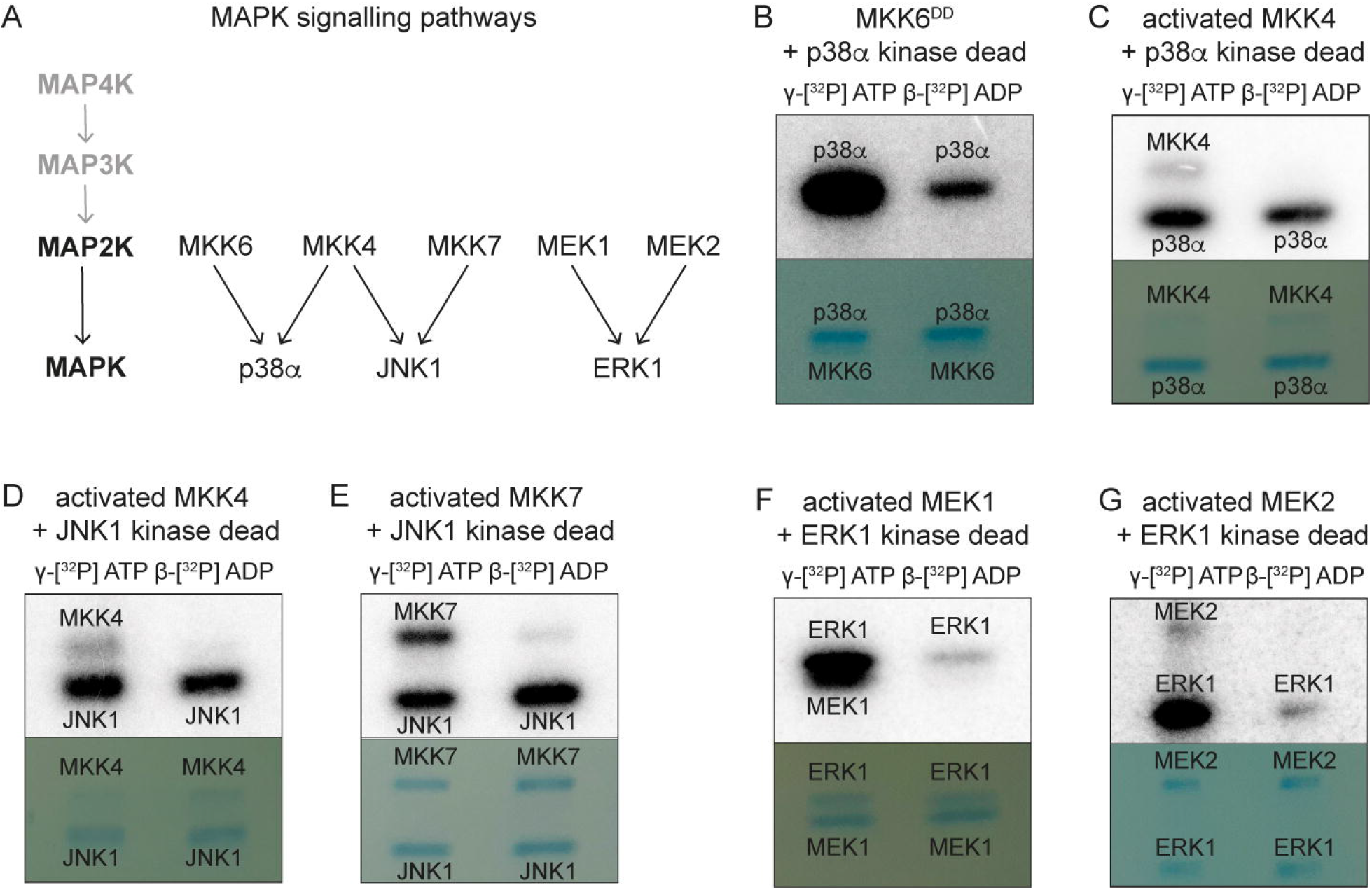
All MAP2Ks can use ADP to phosphorylate their MAPK substrates. *In vitro* phosphorylation assay with radiolabelled nucleotides. Pairs of activated MAP2K (MKK6^DD^, MKK4, MKK7, MEK1 and MEK2) and kinase dead MAPK (p38α, JNK1 and ERK1) were incubated with γ-[^32^P] ATP or β-[^32^P] ADP. Samples were separated by SDS-PAGE and visualized by autoradiography (top) and Coomassie stain (bottom).

As a further control, we also tested the ability of a kinase from a different branch (TLK) of the kinome(22), the kinase domain of RIPK2 (23), to use ADP as a phosphate donor (Figure 4). RIPK2 activation occurs through autophosphorylation at the A-loop, where up to 6 closely located phosphorylation sites are present (23, 24). Moreover, during signalling, RIPK2 can phosphorylate a tyrosine residue in the RIPK2 CARD domain (25), showing the ability of this kinase to phosphorylate both serine/threonine and tyrosine residues, unusual in protein kinases, making it similar to MAP2Ks. Our data demonstrate that this kinase is not able to use ADP to autophosphorylate, suggesting the use of ADP as substrate is not a universal mechanism in protein kinases. However, it remains possible that other kinases more closely related to the MAP2Ks could have this ability as well.

**Figure 4.**
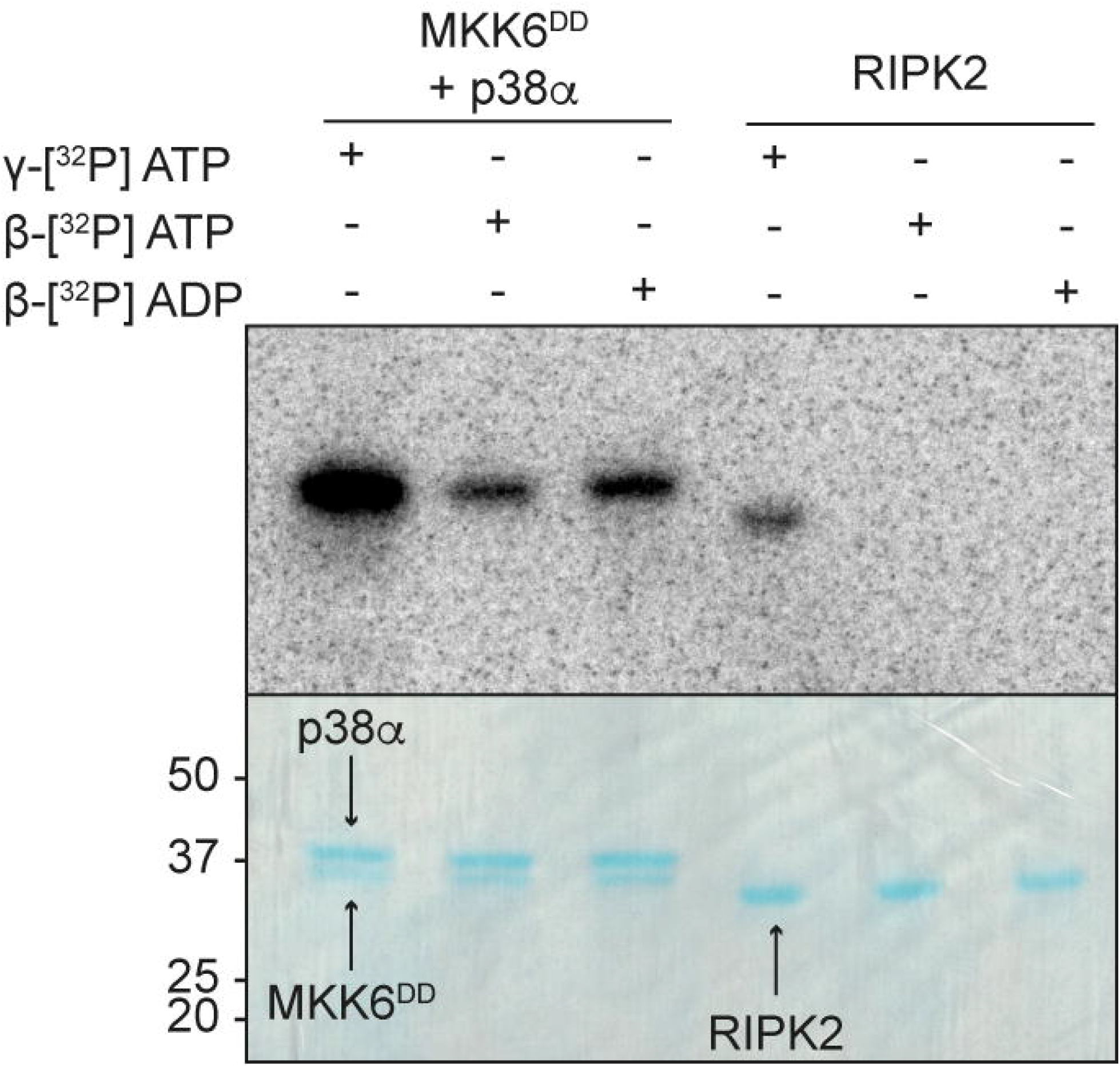
RIPK2 cannot use ADP to autophosphorylate. *In vitro* phosphorylation assay with radiolabelled nucleotides. Samples of MKK6^DD^ + p38α and RIPK2 were incubated with γ-[^32^P] ATP, β-[^32^P] ATP, or β-[^32^P] ADP. Samples were separated by SDS-PAGE and visualized by autoradiography (top) and Coomassie stain (bottom).

## Discussion

The majority of phosphoryl transfer enzymes are dependent on ATP as the source of phosphate. A small number of kinases have evolved to position ADP for nucleophilic attack on the β-phosphate. These ADP dependent kinases were initially discovered in extremophiles where it was thought that high temperatures could lead to depleted levels of ATP. However, subsequent discovery of ADP dependent kinases in mesophiles, and even in vertebrates, has shown that they are more likely adapted for alternative metabolic routes or sensing the state of the cell (26). These ADP dependent kinases are rare and have evolved to use ADP specifically and, to date, no kinases that can use both ATP and ADP have been identified. We have demonstrated that the human dual specificity protein kinases of the MAP kinase pathway, the MAP2Ks, are able to phosphorylate and activate their target MAPKs using both ATP and ADP *in vitro*. The incorporation of the β-phosphate occurs both in the presence and absence of ATP but is less efficient than the use of the γ-phosphate and is not essential to activate the target MAPK. This suggests that the use of ADP is an alternative route to MAPK activation, for example when the concentration of ATP is very low.

MAP2Ks are an unusual family of protein kinases referred as “dual specificity kinases”: they can phosphorylate two chemically and structurally different amino acids, tyrosine and threonine, separated by a single residue (TxY motif). The majority of protein kinase families phosphorylate either the structurally similar residues serine and threonine, or are specific for tyrosine (and in some cases histidine). Recent structural studies from our laboratory have demonstrated that, as opposed to a classical enzyme, where substrates are precisely positioned, the MAP2K MKK6 engages with its substrate MAPK p38α at sites distal from the A-loop (18). This allows flexibility in the amino acid side chain targeted for phosphorylation and could also allow either ATP or ADP to be approached for nucleophilic attack using the same active site architecture. As the β-phosphate is deeper in the active site of the MAP2K, and therefore less accessible to the A-loop, this could account for the lower efficiency observed with ADP phosphorylation of p38α. In addition to the lower efficiency of activation there is also a lower proportion of doubly phosphorylated p38α. This could allow for alternative signalling, as it has been suggested that monophosphorylated p38α and ERK could have alternative signalling roles (27-29).

As the MAPK pathways are generally involved in the stress response, it is conceivable that the ability to maintain signalling in a low ATP environment would be an advantage, for instance under hypoxic conditions, that occur under a number of cell stresses, such as cancer and atherosclerosis. It has been clearly demonstrated that MAP kinase signalling is maintained, and often activated, during these hypoxic conditions (30-33). Cells are extremely sensitive to the ATP/ADP ratio with many proteins sensitive to the ratio triggering metabolic responses, maintaining stress signalling under these circumstances is also important. The ATP/ADP ratio drops significantly within minutes of ischemia, for example from 10 to 0.4 within 90 seconds in rat cerebellum (34). The use of ATP in the cell depends on its displacement from equilibrium, that is lost upon the collapse of the ATP/ADP ratio, a problem that could be circumvented by the alternative use of ADP. The observation that the MAP2Ks can use ADP to activate their target MAPKs *in vitro* is surprising and needs to be demonstrated if and when it occurs *in vivo*. The observation also means that kinase kinetic assays based on the formation of ADP should be reassessed. The phenomenon may be limited to the unusual dual specificity MAP2Ks, but it would seem prudent to test as wide a range of protein kinases as possible in order to determine how limited the ability to use ADP is in protein kinases. This study is an invitation to all kinase researchers to test their kinase’s ability to phosphorylate using ADP.

## Methods

### Plasmids

Plasmids for protein expression were ordered from Genscript (gene synthesis, cloning and mutagenesis). The sequences for p38α, ERK1 and JNK1 (WT and kinase dead mutants) were fused to a His6 tag and 3C protease cleavage site and cloned into pET-28b vector. MKK6 sequences (WT and constitutively active S207D T211D mutant, referred to as MKK6^DD^) were fused to a twin StrepII tag and 3C cleavage site and cloned into a pFastBac1 vector. Lambda Phosphatase plasmid was ordered from Addgene (a gift from John Chodera & Nicholas Levinson & Markus Seeliger; Addgene plasmid #79748; http://n2t.net/addgene:79748; RRID:Addgene_79748) (35).

### Protein expression and purification

Constructs of p38α, ERK1 and JNK1 were co-transformed with lambda phosphatase plasmid into Rosetta™(DE3)pLysS *E. coli* competent cells (Novagen) with appropriate antibiotics. Cells were grown in LB at 37°C until the OD600 = 0.6-0.8, induced with 0.5 mM IPTG, incubated at 16°C overnight, and harvested by centrifugation. MKK6 constructs were transformed into DH10 *E. coli* cells to produce recombinant baculoviruses subsequently used for protein expression in *Sf21* insect cells (36). Cells were harvested 48h after proliferation arrest by centrifugation.

Cell pellets were resuspended in lysis buffer (50 mM HEPES pH 7.5, 200 mM NaCl, 10 mM MgCl_2_, 5% glycerol, 0.5 mM TCEP, with a Pierce protease inhibitor EDTA-free tablet (Thermo Scientific) and traces of DNaseI (Sigma)). Cells were lysed by sonication on ice and the lysate centrifugated. For p38α, ERK1 and JNK1, supernatant was loaded onto a pre-packed 5 ml HisTrap column (GE Healthcare), equilibrated according to the supplier’s protocols with wash buffer (50 mM HEPES pH 7.5, 200 mM NaCl, 10 mM MgCl_2_, 5 % glycerol, 0.5 mM TCEP) with 1% of elution buffer (wash buffer with 500 mM Imidazole). Tagged-protein was eluted with elution buffer, and target protein-containing fractions were pooled, incubated with 3C-protease and dialysed against wash buffer at 4°C overnight. The sample was run through the HisTrap column again and flow-through fractions were collected. For MKK6, supernatant was loaded onto a pre-packed 5 ml StrepTactin XT column (IBA), equilibrated according to the supplier’s protocols with wash buffer (50 mM HEPES pH 7.5, 200 mM NaCl, 10 mM MgCl_2_, 5 % glycerol, 0.5 mM TCEP). Tagged-protein was eluted with 50 ml elution buffer (wash buffer with 50 mM Biotin), and MKK6-containing fractions were pooled, incubated with 3C-protease and dialysed against wash buffer at 4 °C overnight. The sample was run through the StrepTactin XT column again and flow-through fractions were collected.

The RIPK2 kinase domain was expressed and purified as previously described (23). Pure proteins were concentrated using centrifugal filter units (Amicon Ultra, 10 kDa MWCO), flash-frozen in liquid nitrogen and stored at −80°C before being used for biochemical assays.

Activated recombinant MAP2K proteins were purchased from Proquinase (MEK1, MKK3, MKK4 and MKK7), and R&D Systems (MEK2).

### Nucleotides

Nucleotides and analogues were purchased from Merck, ultra-pure ADP was purchased from Cell Technology.

### Phosphorylation assays using native PAGE gels

For end point phosphorylation assays, samples were prepared on ice in 25 μl reaction volumes with 1 μM MKK6^DD^ and 1 μM p38α in assay buffer (50 mM HEPES pH 7.5, 200 mM NaCl, 10 mM MgCl_2_, 5 % glycerol, 0.5 mM TCEP). Nucleotides were added to a final concentration of 0.5 mM (unless specified), and samples were incubated for 20 min at 30°C. For the inhibitor assay, 0.5 mM Staurosporine (Merck, 569397) was added prior to the nucleotide. Native sample buffer (final concentrations: 10% glycerol, 20 mM Tris pH 6.8, traces of bromophenol blue, 25 mM EDTA) was added to quench the reactions.

For time course phosphorylation assays, samples were prepared in 500 μl reaction volumes with 0.2 μM MKK6^DD^ and 1 μM p38α in assay buffer (50 mM HEPES pH 7.5, 200 mM NaCl, 10 mM MgCl_2_, 5 % glycerol, 0.5 mM TCEP). Nucleotides were added to a final concentration of 1 mM, and time measurements started. At given time points, samples were taken and quenched in native sample buffer (final concentrations: 10% glycerol, 20 mM Tris pH 6.8, traces of bromophenol blue, 25 mM EDTA). Samples were loaded on Pre-cast 4-20% gradient Tris-Glycine gels (ThermoFisher Scientific) and the gel run in Tris-Glycine Native running buffer (2.5 mM Tris Base, 19.2 mM glycine pH 8.3) at 125 V for 1h30 min. Gels were stained with InstantBlue (Expedeon) or Coomassie staining (when for MS analysis), imaged with a ChemiDoc system (Bio-Rad) and analysed in ImageLab software (v6.0, Bio-Rad).

### Mass spectrometry

Protein samples were prepared as for the native-PAGE gel analysis, flash-frozen in liquid nitrogen and stored at −80°C until analysis.

#### Intact mass by Q-TOF MS (Quadrupole time-of-flight Mass Spectrometry)

Protein samples were acidified using 1% TFA prior to the injection of ~5 μg of each sample onto an Acquity UPLC Protein BEH C4 column on the Acquity UPLC System (Waters Corporation) coupled to a quadrupole time of flight (Q-TOF) Premier mass spectrometer (Waters/Micromass) using the standard ESI source in positive ion mode. Solvent A was water, 0.1% formic acid, and solvent B was acetonitrile, 0.1% formic acid. Data was acquired in continuum mode over the mass range 500–3500 m/z with a scan time of 0.5 s and an interscan delay of 0.1 s. Data were externally calibrated against a reference standard of intact myoglobin, acquired immediately prior to sample data acquisition. Spectra from the chromatogram protein peak were then summed and intact mass was calculated using the MaxEnt1 maximum entropy algorithm (Waters/Micromass) to give the zero charge deconvoluted molecular weight.

#### Digestion and PTMs analysis by LC-MS/MS (Liquid chromatography coupled to tandem Mass Spectrometry)

Samples were prepared following the SP3 protocol (37). Analysis was performed on an UltiMate 3000 RSLC nano LC system (Dionex) fitted with a trapping cartridge (μ-Precolumn C18 PepMap 100) and an analytical column (nanoEase™ M/Z HSS T3 column, Waters) coupled directly to a Fusion Lumos (Thermo) mass spectrometer using the proxeon nanoflow source in positive ion mode.

The peptides were introduced into the Fusion Lumos via a Pico-Tip Emitter (New Objective) and an applied spray voltage of 2.4 kV. Full mass scan was acquired with mass range 375-1200 m/z in profile mode in the orbitrap with resolution of 120000. The filling time was set at maximum of 50 ms with a limitation of 4×105 ions. Data dependent acquisition (DDA) was performed with the resolution of the Orbitrap set to 30000, with a fill time of 86 ms and a limitation of 2×105 ions. A normalized collision energy of 34 was applied. MS2 data was acquired in profile mode. Acquired data were processed by IsobarQuant (38), the Mascot (v2.2.07) search engine was used.

### Phosphorylation assays using radiolabelled nucleotides

Protein samples were prepared on ice at 0.5 μM in 10 μl assay buffer (50 mM HEPES pH 7.5, 200 mM NaCl, 10 mM MgCl_2_, 5 % glycerol, 0.5 mM TCEP). Radiolabelled nucleotide sources (γ-^32^[P] ATP, β-[^32^P] ATP, and β-[^32^P] ADP (Hartmann Analytic GmbH)) were diluted 1:10 into a 1 mM cold nucleotide solution (ATP or ADP as appropriate). The reaction was set up by adding 1 μl of nucleotide to each sample and incubated for 20 min at 30°C. The reaction was stopped by adding 4 μl of SDS sample buffer (0.4% bromophenol blue, 0.4 M DTT, 0.2 M Tris pH 6.8, 8% SDS, 40% glycerol) and boiling for 5 min at 95 °C. Samples were centrifuged and loaded on Pre-cast 4-20% gradient Tris-Glycine gels (ThermoFisher Scientific), run in Tris-Glycine running buffer (2.5 mM Tris Base, 19.2mM glycine pH 8.3, 1% SDS) at 220 V for 40 minutes. The gels were exposed to a storage phosphor screens (GE) overnight and imaged using a Typhoon scanner (GE Health). Gels were then stained with InstantBlue (Expedeon) and imaged. Images were analysed using Image Lab (Bio-Rad) software.

### ^31^P NMR spectra of nucleotides stocks

^31^P NMR spectra of nucleotides (ATP or ADP) at 10 mM in buffer were acquired on a Bruker 700 MHz spectrometer equipped with a cryoprobe at 25°C and processed with TopSpin. Peaks were assigned by comparison with published work (39).

## Acknowledgments

We thank A. Aubert and M. Pelosse (EMBL Grenoble) for support in eukaryotic protein expression, O. Pessey (EMBL Grenoble) for support with radiolabelling assays, Mandy Rettel and Dominic Helm at the proteomic core facility (EMBL Heidelberg) for mass spectrometry, and Malene Jensen and Elise Delaforge (IBS Grenoble) for assistance with NMR experiments.

## Author contributions

Conceptualization, E.P and M.W.B; methodology, P.J, J.vV and E.P,; investigation, P.J, J.vV, E.P and M.W.B; visualization, P.J, J.vV; funding acquisition, M.W.B; project administration, M.W.B; supervision, E.P and M.W.B; writing – original draft, P.J, E.P and M.W.B; writing – review & editing, P.J, J.vV, E.P and M.W.B.

## Figure legends

**Figure S1.**
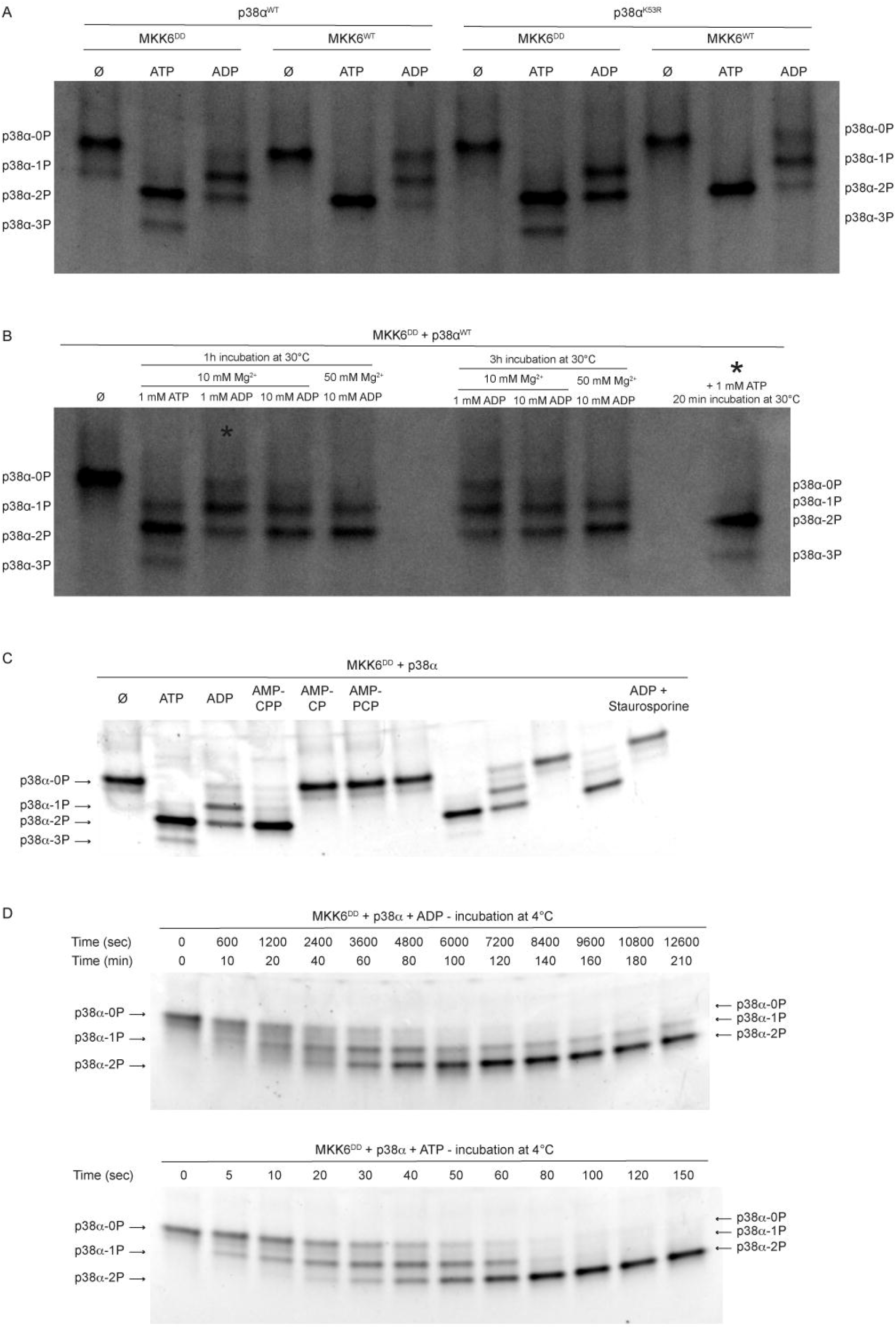
ADP phosphorylation of p38α variants by MKK6 variants under different conditions. (A) Native PAGE gel of MKK6 (WT or constitutively active DD mutant) + p38α (WT or kinase-dead K53R mutant) in the presence of nucleotide. p38α bands run based on the total phosphorylation number of the protein. (B) Native PAGE gel of MKK6^DD^ + p38α^WT^ in the presence of different nucleotides and Mg^2+^ concentrations, and longer incubation time. The last sample was initially incubated for 1h at 30°C with 1 mM ADP, and was then supplemented with 1 mM ATP and incubated for 20 additional minutes before being run on the gel. p38α bands run based on the total phosphorylation number of the protein. (C) Uncropped gel from Fig 1A. (D) Representative gels used to generate Fig 2.

**Figure S2.**
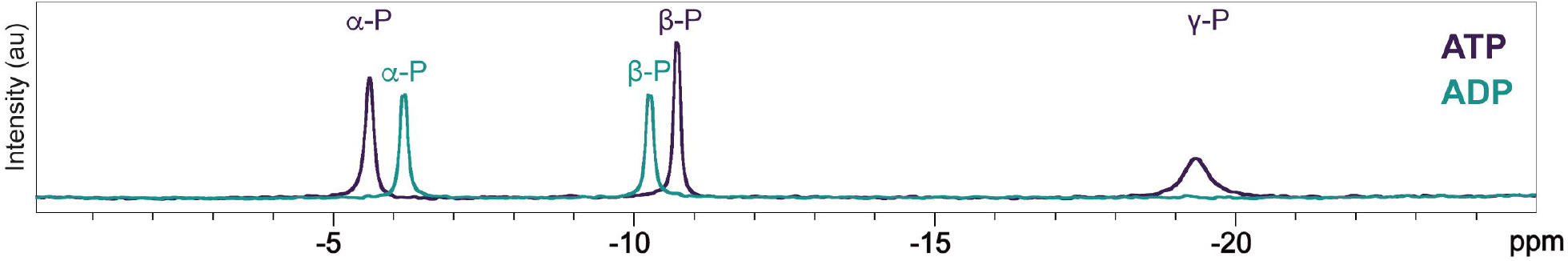
Nucleotide purity. Natural abundance ^31^P NMR spectra of ATP and ADP stock solutions used in *in vitro* assays demonstrating the absence of ATP in the ADP solutions.

